# Interpretable Distillation Reveals that Deep-learning-based Splicing Models Suffer from Pervasive Confounders and Blind Spots

**DOI:** 10.1101/2025.11.28.691252

**Authors:** Simon Liu, Wenjing Zhang, Oded Regev

**Affiliations:** New York University, New York, NY, 10012, USA

## Abstract

Despite their growing popularity, genomic deep-learning-based models function largely as black boxes, raising concerns about their trustworthiness. Here we develop a framework to explain model prediction logic using interpretable distillation. Applying our framework, we find that RNA splicing prediction models suffer from pervasive confounders and blind spots, leading to poor performance on non-reference sequences. Our findings illuminate fundamental limitations of training models on genomic sequences and suggest ways to overcome them.

## INTRODUCTION

Deep learning is rapidly emerging as a central approach for deciphering how genomic sequences encode gene regulatory information.^1,2^ But despite their growing use, current models depend on complex, black-box-like architectures,^3–5^ leaving their predictive logic poorly understood and raising concerns about their trustworthiness.^6^ Prior methods to interpret models have primarily focused on local interpretation, explaining individual predictions rather than overall model logic.^3–5^ Methods are therefore needed to interrogate the global logic of genomic deep-learning-based models, assess their reliability across diverse contexts, and identify conditions under which they fail.

Here, we present an interpretable distillation framework to elucidate the logic of genomic deep-learning-based models and demonstrate its applicability to RNA splicing prediction models,^4,5,7^ which are being increasingly deployed in clinical settings.^8,9^ With this approach, we train an interpretable, “distilled” model that closely replicates the predictions of a deep-learning-based model and then analyze it as a proxy to infer the original model’s logic.^10,11^ Using our framework, we find that splicing models recognize exons through surprisingly simple additive combinations of sequence motifs, including known splicing regulatory elements. Critically, our analysis also reveals that splicing models exploit genomic confounders unrelated to splicing and fail to capture RNA structure effects, leading to systematic prediction errors. Our findings highlight fundamental limitations of genomic deep learning that extend beyond splicing prediction and underscore the importance of model interpretation for assessing trustworthiness.

## RESULTS

### Splicing Models Recognize Exons through Additive Combinations of Motifs

To interpret deep-learning-based splicing models, we trained distilled models to replicate SpliceAI,^4^ Pangolin,^7^ and AlphaGenome^5^ predictions on a certain exon recognition task (**Figure 1A**). Our models’ predictions are formed by simply summing individual contributions from sequence features, enabling complete interpretability.

**Figure 1.**
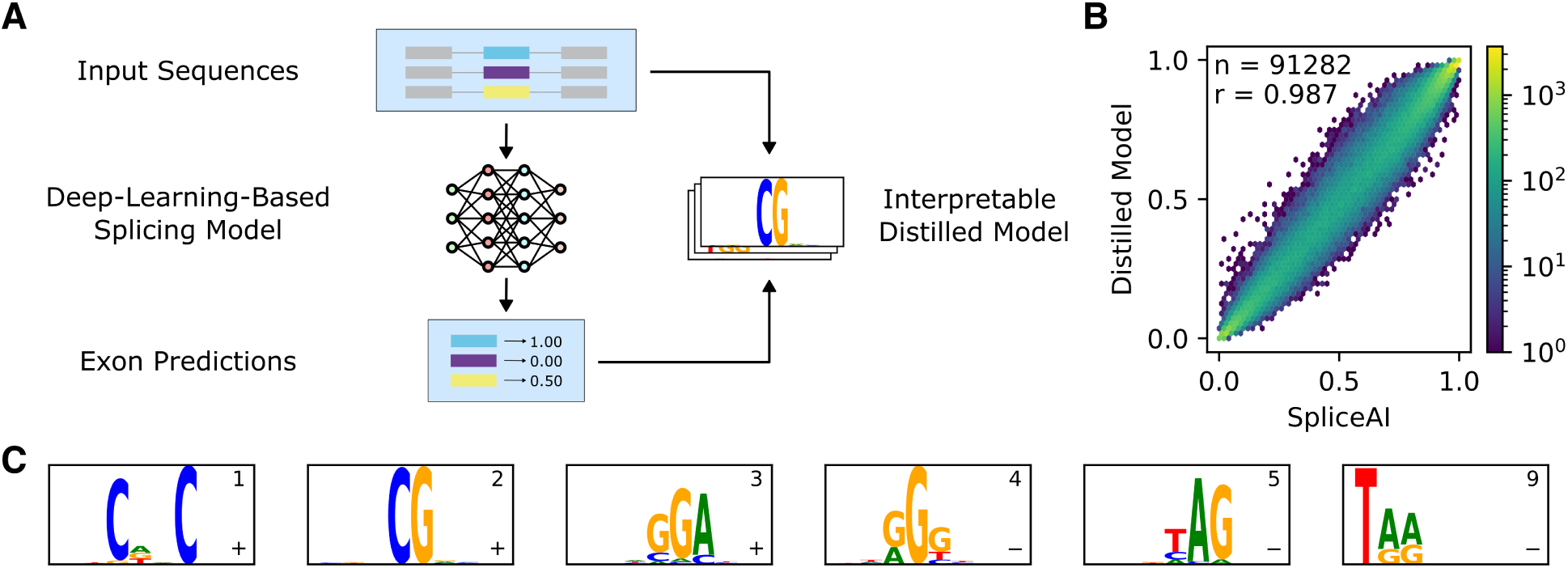
Interpretable distillation reveals exon recognition logic of deep-learning-based splicing models. **(A)** Interpretable distillation framework. An interpretable model is trained to replicate exon predictions of a deep-learning-based splicing model. **(B)** Comparison of distilled model predictions to SpliceAI predictions on held-out sequences. **(C)** Selection of sequence features, sorted by importance rank (top right number), learned by distilled model of SpliceAI. Sequence logos show convolutional kernel preferences. Effect directionality shown in bottom right (+: increase; –: decrease). See **Figure S1A** for all features learned from SpliceAI, **Figure S2** for Pangolin, and **Figure S3** for Al-phaGenome.

We found that the distilled models replicate each splicing model’s predictions almost perfectly (Pearson *r* : 0.94 – 0.99 across models on held-out data) (**Figures 1B**, **S2**, **S3A**), validating them as reliable proxies for interpretation. For SpliceAI, 98.4% of the distilled model’s predictions fall within *±*15% of the original’s model prediction. Remarkably, the distilled models achieve this accuracy without modeling RNA structure or feature interactions, indicating that deep-learning-based splicing models recognize exons primarily through simple additive sequence feature combinations.

By inspecting the sequence features learned by the distilled models, we found that splicing models identify motifs resembling splicing regulatory elements^12–15^ (**Figures 1C**, **S1A**, **S2**, **S3B**). Notably, we also discovered a motif that matches a liver-specific^16^ regulatory element in that tissue’s Pangolin model, illustrating how interpretable distillation captures a broad range of predictive features (**Figure S2C**). Furthermore, we identified several features not immediately matching known RNA-binding protein preferences, potentially representing uncharacterized regulatory elements.^12,13^ Together, these findings demonstrate that interpretable distillation effectively decodes the predictive logic of deep-learning-based splicing models.

### Splicing Models Confound Coding Sequence CpG Enrichment with Exon Identity

Across all models, CG dinucleotides (CpG) consistently emerged as one of the two most influential features for distinguishing exons (**Figures 1C**, **S2**, **S3B**). Surprisingly, CpG does not match any known splicing factor binding motif and is shorter than typical RNA-binding protein binding sites.^12,13^ This suggests that CpG is unlikely to directly regulate splicing. Moreover, we considered the possibility that CpG regulates splicing indirectly through genomic methylation; however, we failed to find any association between genomic CpG methylation and exon inclusion (**Fig-ure S4**),^17^ suggesting that, in general, CpG methylation does not noticeably affect splicing. In summary, these observations indicate that CpG represents a confounding feature.

To quantify how strongly CpG confounds splicing models, we compared model predictions to experimentally measured exon inclusion levels, defined as the percent of transcripts with a given exon (percent spliced in; PSI), on three experimental assays covering diverse sequence contexts.^18–20^ Even though all three models generally provide accurate predictions for all three assays (Pearson r for Liao et al. 2023: 0.78 – 0.85, *FAS* exon 6: 0.75 – 0.84, MFASS: 0.40 – 0.45), we found that CpG composition is significantly associated with prediction error (ANCOVA, p < 5 *×* 10^−3^ in all cases) (**Figures 2A**, **S5**). Namely, we observed that sequences with higher CpG content consistently received inflated predictions, while sequences with lower CpG content received deflated predictions, even when controlling for measured inclusion levels. We did not observe a consistent association between GpC composition and prediction error, indicating that this impaired reliability is specific to CpG rather than overall GC content (**Figure S6**).

**Figure 2.**
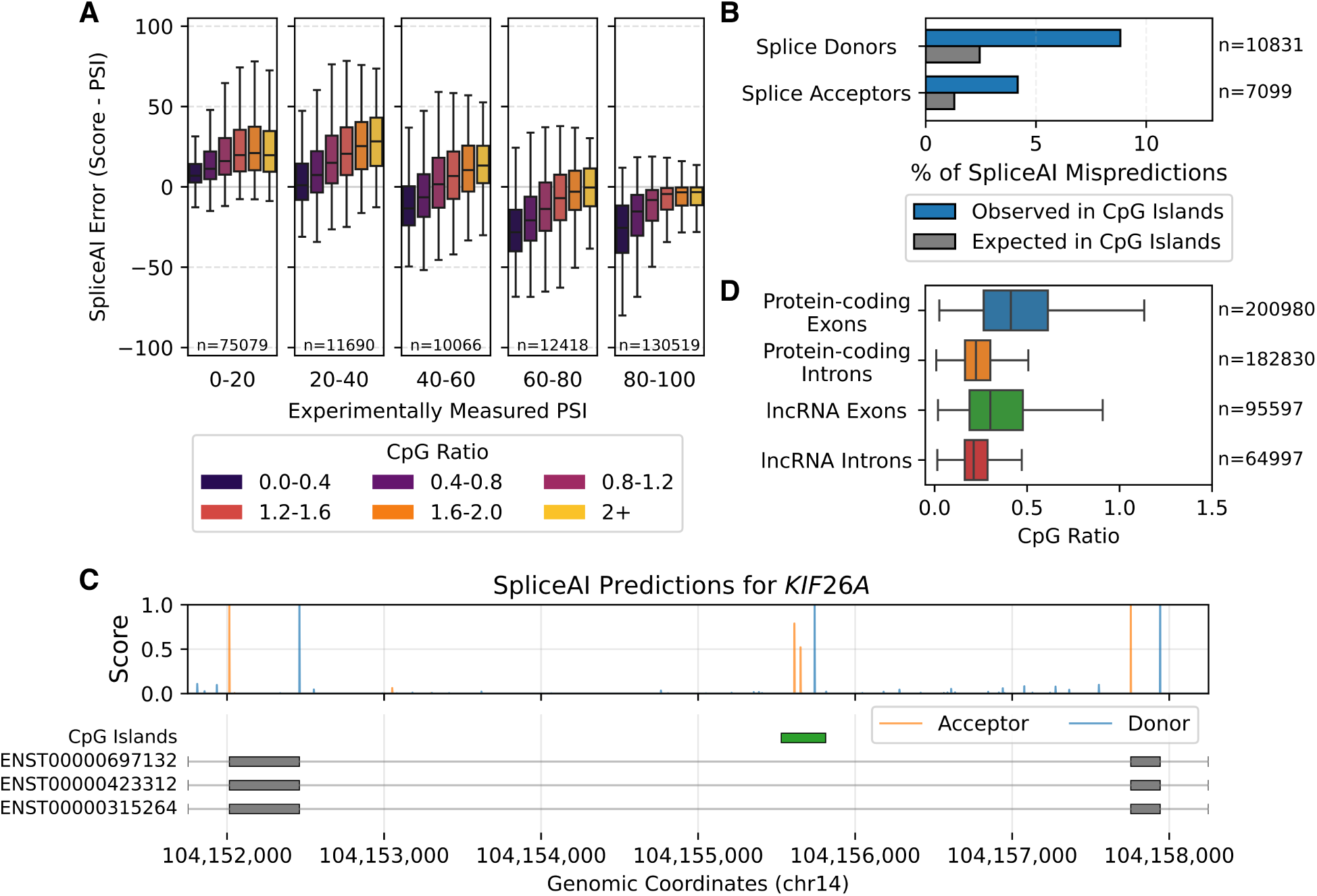
Deep-learning-based splicing models learn CpG enrichment as a spurious marker of exons. **(A)** SpliceAI error (prediction minus measured inclusion) on an experimental splicing assay.^18^ Box plots show prediction error across CpG ratios (defined as the observed frequency of CG dinucleotides divided by the expected frequency based on overall GC content). Each box plot contains *≥* 150 sequences. See **Figure S5** for analysis with other splicing models and assays. **(B)** Percentage of false positive SpliceAI predictions (score > 0.2, unannotated sites) within intronic CpG islands compared to matched control regions. **(C)** SpliceAI splice site predictions in *KIF26A*, with CpG islands and Ensembl transcript annotations. **(D)** CpG ratio in protein-coding and lncRNA gene regions. Boxes in (A) and (D) show quartiles, with whiskers extending to the full distribution excluding outliers.

Next, we examined the impact of CpG bias on genomic predictions. We found that SpliceAI’s false positive predictions (score > 0.2) cluster near intronic CpG islands, particularly for splice donors (**Figure 2B**). For example, SpliceAI predicts a strong donor-acceptor pair within an intronic CpG island in *KIF26A*, despite the absence of annotated splice sites (**Figure 2C**). Overall, we found that excluding false predictions within CpG islands improves SpliceAI’s splice donor prediction top-k accuracy for genes with CpG islands from 93.8% to 94.5% (11.3% error reduction), demonstrating that CpG bias is a major source of false genomic predictions.

Why do splicing models learn this spurious CpG bias? Genomic protein-coding exons are enriched in CpG relative to introns (**Figure 2D**), an enrichment that likely reflects properties of coding sequences rather than splicing regulation. For instance, long non-coding RNA (lncRNA) transcripts, which undergo splicing but lack coding constraints, show reduced exonic CpG enrichment (**Figure 2D**). Consistent with its reliance on CpG, SpliceAI’s performance drops markedly on lncRNA (84% top-k accuracy) compared to protein-coding transcripts (95% top-k accuracy).^4^ We hypothesize that splicing models, having been trained on protein-coding genes, exploit CpG enrichment as a proxy for exon identity despite lacking a mechanistic connection to splicing.

### Splicing Models Confound Stop Codon Depletion with Exon Identity

By examining other influential features, we also discovered that stop codons (UAA, UAG, UGA) consistently lower predictions across all models (**Figures 1C**, **S2**, **S3B**). Their effect exhibits a three-nucleotide periodicity (**Figures S1B**, **S2**, **S3B**), further suggesting that splicing models recognize translational constraints. However, current evidence does not support any direct role of stop codons in splicing,^21^ which occurs in the nucleus and precedes translation.

To quantify how strongly stop codons confound splicing models, we again analyzed model predictions compared to measured exon inclusion levels on experimental assay data.^18^ This revealed that stop codons are significantly associated with exon inclusion prediction error for all models (ANCOVA, p < 1 *×* 10^−20^ in all cases) (**Figures 3A**, **S7A**). Specifically, sequences lacking stop codons received inflated predictions, while sequences containing stop codons received deflated predictions, even when controlling for measured inclusion levels. This bias intensified with higher numbers of stop codons and was consistent across all stop codon types (**Figures S7B**, **S7C**, **S7D**). Next, we investigated the effect of stop codon bias on genomic predictions. We found that splicing model predictions for genomic internal protein-coding exons containing stop codons, i.e., at the end of coding regions, are significantly lower than for those without stop codons (**Figures 3B**, **S8A**), even after controlling for exon length and rank (**Figures S8C**, **S8D**). To test whether alternative splicing might explain this difference, we analyzed only constitutive exons and observed the same effect (**Figure S8B**), thus ruling it out as the cause. Critically, this stop codon bias may lead to systematic mispredictions for nonsense variants, which introduce premature stop codons. For example, the *CFTR* G542X variant (c.1624G>T), the most common cystic fibrosis-causing nonsense variant, has been documented to not alter splicing in patient cells or minigene assays,^22–24^ yet both SpliceAI and Pangolin incorrectly predict splice-disrupting effects (**Figure 3C**). This suggests stop codon bias may broadly affect nonsense variant interpretation, undermining clinical reliability.

**Figure 3.**
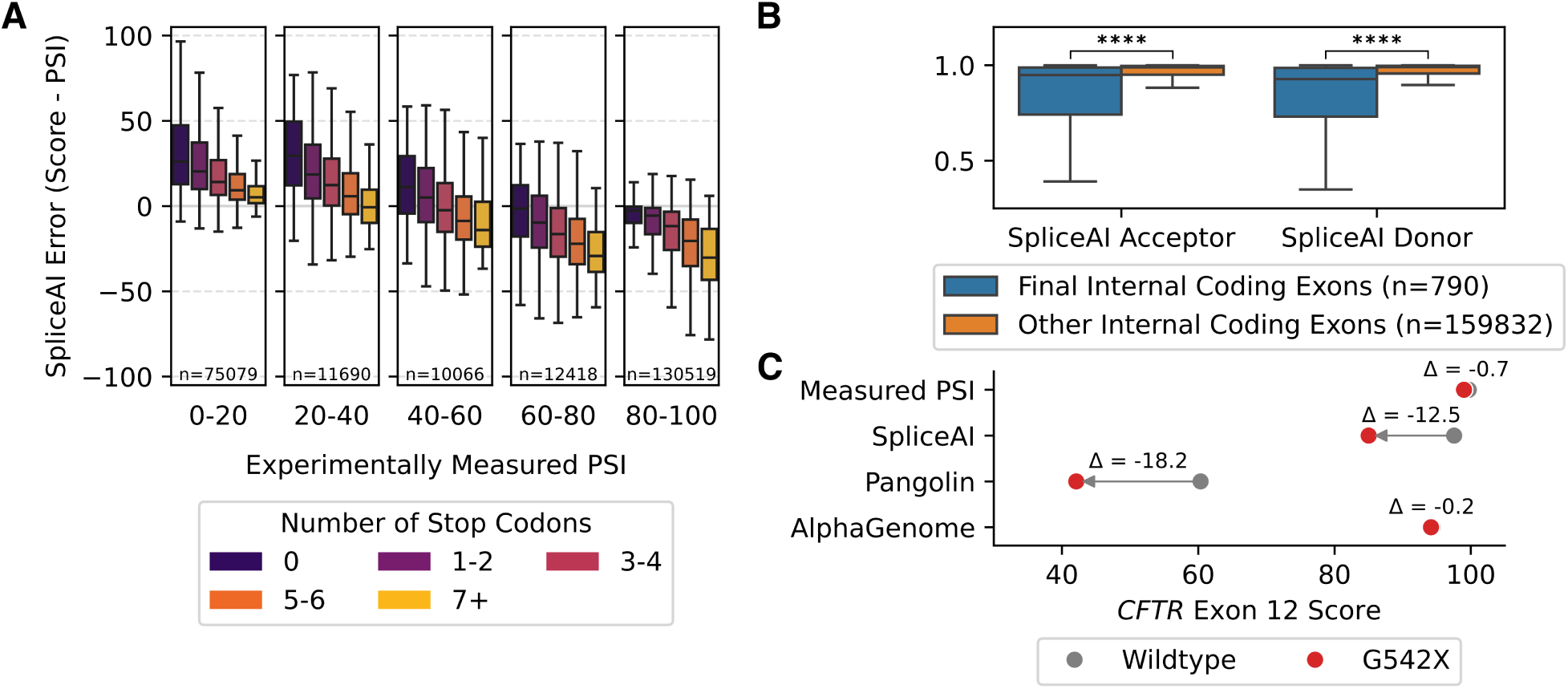
Deep-learning-based splicing models learn stop codon depletion as a spurious marker of exons. **(A)** SpliceAI error (prediction minus measured inclusion) on an experimental splicing assay.^18^ Box plots show prediction error across stop codon count. Each box plot contains *≥* 90 sequences. See **Figure S7** for analysis with other splicing models. **(B)** SpliceAI acceptor and donor scores for genomic internal coding exons in MANE Select transcripts. Two-tailed Welch’s t-test: ****: P *≤* 0.0001. **(C)** Measured PSI^22^ and model predictions for *CFTR* exon 12 wildtype and G542X variant. Boxes in (A) and (B) show quartiles; whiskers extend to the full distribution excluding outliers.

Why do splicing models learn this spurious stop codon bias? In protein-coding genes, stop codons typically appear only in the final coding exon, meaning that exons are generally depleted of stop codons genome-wide. We hypothesize that splicing models exploit this genomic pattern as a proxy for exon identity despite its lack of mechanistic connection to splicing, causing poor predictions for sequences containing stop codons.

### Splicing Models Fail to Learn the Effects of RNA Structure

RNA structure has been widely established as a modulator of exon recognition,^18,25–27^ and previous work has demonstrated that incorporating structural features significantly improves splicing prediction accuracy.^18,27^ However, our distilled models reveal that splicing models lack structure-dependent logic for recognizing exons (**Figures S1A**, **S2**, **S3B**).

To measure how this “blind spot” affects prediction accuracy, we again compared model predictions to measured exon inclusion levels on assay data.^18^ We found that models consistently inflate predictions for synthetic sequences with more thermodynamically stable predicted structures (ANCOVA, p < 1 *×* 10^−20^ in all cases) (**Figures 4A**, **S9A**). To further validate this finding, we analyzed predictions for *MAPT* exon 10 variants with structures experimentally characterized by chemical probing.^27^ We similarly observed that variants that increase structural stability are associated with inflated predictions (**Figures 4B**, **S9B**), demonstrating that failure to model RNA structure effects contributes to a substantial portion of prediction error.

**Figure 4.**
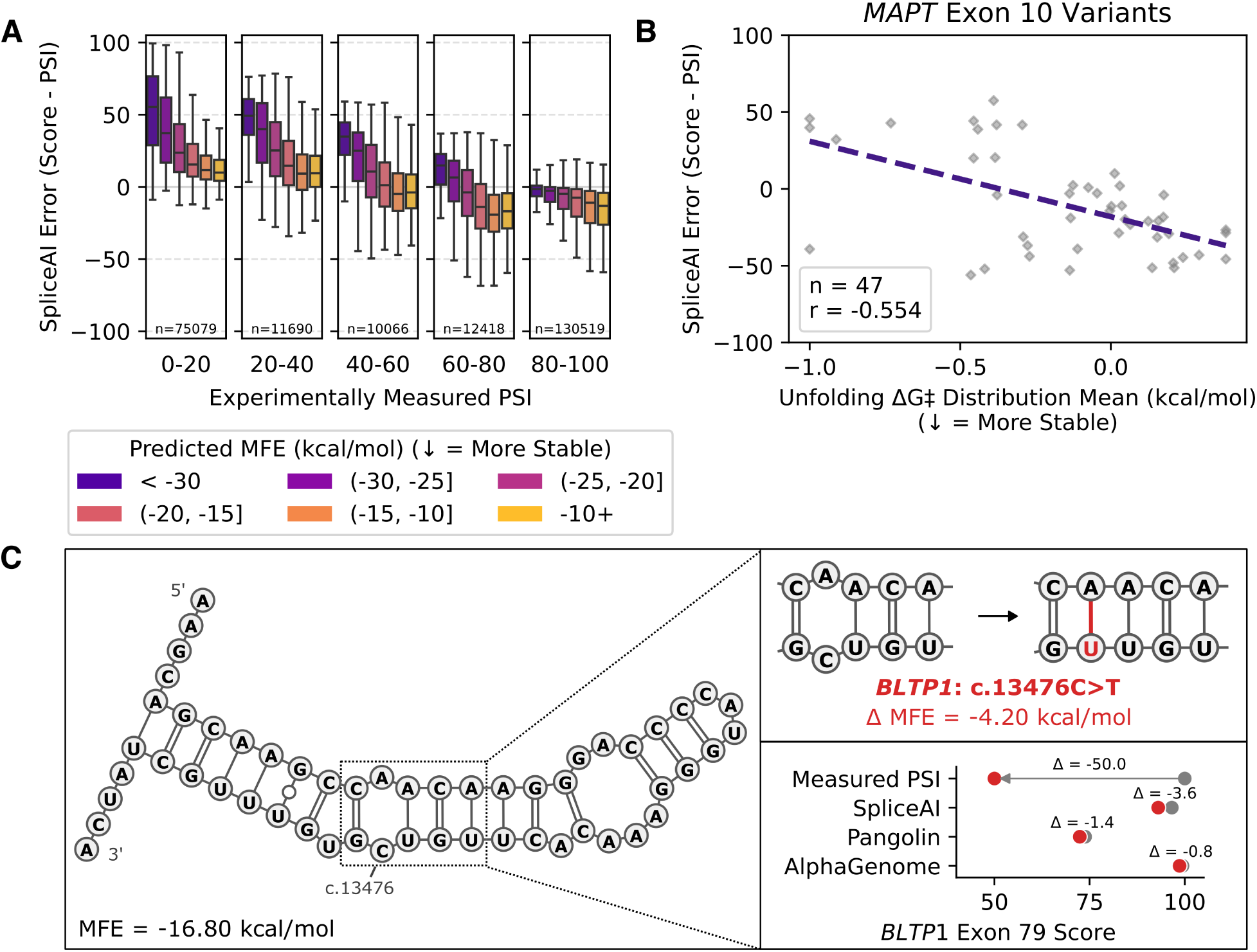
Deep-learning-based splicing models fail to learn RNA structure effects. **(A)** SpliceAI error (prediction minus measured inclusion) on an experimental splicing assay.^18^ Box plots show prediction error across predicted minimum free energy (MFE). Boxes show quartiles; whiskers extend to the full distribution excluding outliers. Each box plot contains *≥* 90 sequences. See **Figure S9A** for analysis with other splicing models. SpliceAI error as a function of RNA structure score for *MAPT* exon 10 variants with experimentally characterized structures. See **Figure S9B** for analysis with other splicing models. **(C)** Predicted RNA secondary structures for *BLTP1* exon 79 wildtype (left) and *BLTP1*:c.13476C>T variant (top right), with splicing model predictions (bottom right).

This blind spot has critical implications for variant interpretation. For example, the *BLTP1* c.13476C>T variant, which causes epileptic encephalopathy, was recently highlighted in a case study illustrating the diagnostic limitations of *in silico* splicing prediction tools.^9^ This variant causes exon skipping in patient cells and minigene assays and is predicted to strengthen a stem loop within *BLTP1* exon 79 (**Figure 4C**). Yet SpliceAI, Pangolin, and AlphaGenome all fail to predict any effect on splicing, demonstrating a gap in current models.

Why do splicing models fail to learn the effects of RNA structure? One possibility is that genomic sequences offer limited structural diversity,^28^ leaving splicing models with insufficient examples from which to learn structural principles. Furthermore, accurate RNA structure prediction typically depends on specialized model architectures^29^ or prior knowledge of thermodynamic principles,^30^ which current splicing models do not incorporate. We hypothesize that, together, these constraints create the blind spot to RNA structure observed in splicing models.

## DISCUSSION

We developed an interpretable distillation framework to explain the logic of genomic deep-learning-based models. Unlike *in silico* mutagenesis and other local interpretation methods which explain individual predictions, interpretable distillation provides a global view of model reasoning.^3,10,11^ Applying this approach, we extracted the logic used by SpliceAI, Pangolin, and AlphaGenome to recognize exons within a fixed sequence context. Our analysis revealed fundamental limitations in what models learn, including confounders and blind spots that compromise prediction reliability. Previous efforts to improve splicing prediction have focused on either using larger and more diverse genomic datasets or developing more complex model architectures.^5,7,31^ Yet the persistence of these issues across scale, from convolutional networks such as SpliceAI to foundation models such as AlphaGenome, suggests that they are inherent to the genome-based training paradigm. Genomes contain pervasive correlations that confound genuine regulatory signals,^28^ and neither larger genomic datasets nor more complex architectures alone can overcome these biases, whose effects likely extend beyond splicing to other gene regulatory prediction tasks.

Our findings point to several promising directions for addressing these challenges. Previous studies using massively parallel reporter assays have demonstrated that random sequence libraries, which explore broader and more diverse sequence spaces, enable models to capture rich regulatory logic.^18,28^ Integrating such experimental data with genomic datasets during training may help models disentangle genuine regulatory logic from confounders. Moreover, post-processing approaches such as multi-calibration^32^ can mitigate biases in existing models without retraining by adjusting predictions for subgroups with systematic errors. Finally, given the extensive role of structure in regulating RNA processes,^33^ incorporating structural information into models may enable them to capture critical regulatory logic and improve predictive accuracy.^18,27^

As deep learning becomes increasingly central to deciphering gene regulation in research and clinical settings,^8,9^ understanding model logic is essential to avoid inaccurate predictions and misleading biological interpretations. By enabling the interrogation of model reasoning, interpretability frameworks like ours provide a path toward more robust and trustworthy models.

## METHODS

### Synthetic Training Sequences

To train the distilled models, we generated synthetic sequences following a splicing reporter design used in a prior experimental splicing library.^18^ Each synthetic sequence consists of three exons, for which the middle exon contains a 70-nucleotide variable region with fixed 3’ and 5’ splice sites. For the variable region, we randomly sampled nucleotides with equal probability for the four nucleotide bases. We filtered out any sequences with cryptic splice sites predicted inside the variable region (based on the criteria of MaxEntScan v1.1^34^ splice site score greater than half that of either fixed splice site score). After filtering, we retained approximately 1M sequences. Sequences were then scored using each deep-learning-based splicing model to generate the labels used for model distillation (see Splicing Predictions).

### Splicing Predictions

#### SpliceAI

We obtained SpliceAI predictions for all sequences using the published model (downloaded in May 2025 from https://github.com/Illumina/SpliceAI).^4^ For each scored sequence, we provided up to 10,000 nucleotides of surrounding context. We extracted predicted acceptor and donor probabilities (each averaged across the model ensemble) at annotated splice sites and calculated their average as the exon prediction score.

#### Pangolin

We obtained Pangolin predictions for all sequences using the published model (downloaded in May 2025 from https://github.com/tkzeng/Pangolin).^7^ For each scored sequence, we provided up to 10,000 nucleotides of surrounding context. We extracted predicted acceptor and donor usage scores (each averaged across the model ensemble) for each Pangolin tissue (heart, liver, brain, and testis) at annotated splice sites. For distillation, we calculated a tissue-specific exon prediction score as the average of the acceptor and donor usage scores for each given tissue. For all other analyses, we used an average tissue exon prediction score calculated by taking the average of all tissues’ exon prediction scores.

#### AlphaGenome

We obtained AlphaGenome predictions using the API provided by Google DeepMind (accessed in November 2025).^5^ For each sequence, we provided up to 1,048,576 nucleotides of surrounding context. For each scored exon, we calculated the predicted percent spliced in (PSI) using the splice junction predictions in HeLa-S3 cells as

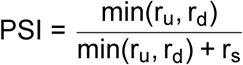

where r_u_ is the predicted junction count between the upstream exon 5’ splice site and the target exon 3’ splice site, r_d_ is the predicted junction count between the target exon 5’ splice site and the downstream exon 3’ splice site, and r_s_ is the predicted junction count between the upstream exon 5’ splice site and the downstream exon 3’ splice site. We removed sequences where the denominator was zero (i.e., no predicted junctions) from final analyses (this only affected the *MAPT* exon 10 variant library^27^). We used predicted PSI as the exon prediction score. For analyses of individual splice site scores, we used the predicted splice site probability tracks. All output from AlphaGenome API is subject to the AlphaGenome Output Terms of Use found at http://deepmind.google.com/science/alphagenome/output-terms.

### Interpretable Distilled Model Architecture

We designed an interpretable neural network architecture that takes as input a one-hot encoded RNA sequence of length L = 120 nucleotides, comprising a 70 nucleotide variable region and 25 nucleotide flanking regions upstream and downstream. First, a convolutional layer with F filters of width k extracts local sequence features. Next, a neural basis layer^35^ assigns position-specific contribution scores to each convolutional filter activation. Finally, contribution scores are summed into a scalar and passed to a multi-layer perceptron to output the final prediction for a sequence. A final sigmoid activation is applied for models trained on KL-divergence loss. L1 regularization was applied during training to convolutional layer activations and neural basis layer activations to promote feature sparsity. This architecture is similar to previous work exploring interpretable machine learning for splicing,^18^ and enables the complete decomposition of predictions into additive contributions from short sequence motifs.

Unlike explanation techniques that yield per-example attributions, our architecture enables global interpretation of the predictive rules learned by a model across an entire dataset. We visualize these rules by representing learned convolutional filters as sequence logos and their corresponding position-specific contribution scores as 1D functions. For each convolutional filter, we scored 10,000,000 random k-mers and computed a weighted position weight matrix for the k-mers corresponding to the top 5% of positive activations. We then visualized sequence logos from these position weight matrices using logomaker.^36^ For the position-specific contribution scores, we evaluated the neural basis layer at each position and filter activation strength to calculate the influence of each convolutional filter when fully activated at a position. We ranked the importance of sequence features by the decrease in Pearson correlation between model predictions and labels on the held-out test set after ablating the convolutional filter.

### Model Distillation Framework

We trained separate distilled models to approximate SpliceAI, Pangolin (one for each of the four tissue models), and AlphaGenome exon predictions. For training, we generated approximately 1M synthetic sequences using a splicing reporter design from prior work (see Synthetic Training Sequences), enabling exploration of a broad space of exonic sequence effects on splicing. We applied each deep-learning-based splicing model to generate exon predictions for these sequences (see Splicing Predictions). We split this dataset into training (*≈*80%), validation (*≈*10%), and test (*≈*10%) sets using random splits.

Distilled models were trained to minimize KL-divergence (for Pangolin, we used mean squared error loss after observing that Pangolin model predictions did not follow a [0, 1] range) between distilled model predictions and target model predictions. Hyperparameter tuning (150 trials) was performed using Bayesian optimization to minimize validation loss. We optimized convolutional kernel number, width, learning rate, batch size, and neural basis model architecture. A full list of hyperparameters and their search ranges is provided in **Table S1**. Models were each trained for up to 200 epochs on NVIDIA A100 GPUs. We implemented our training and evaluation pipeline in PyTorch using Ray^37^ for hyperparameter tuning.

### Experimental Datasets

#### Liao et al. 2023 Synthetic Exon Library

We obtained 239,722 sequences containing random exons with experimentally measured exon inclusion levels from the massively parallel reporter assay published in Liao et al. 2023.^18^ Each synthetic sequence consists of three exons, for which the middle exon contains a 70-nucleotide variable region with fixed 3’ and 5’ splice sites. We downloaded the processed data provided by the authors from https://github.com/regev-lab/interpretable-splicing-model. We used the entire reporter plasmid sequence also provided by the authors to generate splicing predictions.

#### MFASS Library

We obtained 16,469 sequences containing endogenous human exons with experimentally measured exon inclusion levels from the massively parallel splicing reporter assay published in Chong et al. 2019.^20^ We note that this assay uses genomically integrated reporters. For our analysis, we included only wild-type exons and exons with exonic SNVs. We used the original reporter plasmid sequence provided by the authors to generate splicing predictions.

#### *FAS* Exon 6 Mutagenesis Library

We obtained 7,918 sequences (5,976 unique sequences) consisting of *FAS* exon 6 variants with experimentally measured exon inclusion levels from the mutagenesis study published in Baeza-Centurion et al. 2025.^19^ Each sequence represents a SNV, 1- to 4- nucleotide insertion, or deletion from the wild-type *FAS* exon 6. We used the surrounding sequence context from the reference genome for *FAS* to generate splicing predictions.

#### *MAPT* Exon 10 Variant Library

We obtained 47 *MAPT* exon 10 variants with experimentally measured exon inclusion levels reported by Kumar et al. 2022.^27^ Each variant has an experimentally determined structural stability score. We used the surrounding sequence context from the reference genome for *MAPT* to generate splicing predictions. We excluded 4 variants from AlphaGenome-specific analyses as AlphaGenome failed to predict any splice junctions.

### CpG Composition Analysis

For each sequence, we measured CpG composition by CpG ratio, defined as the observed frequency of CG dinucleotides divided by the expected frequency based on overall guanine and cy-tosine content:

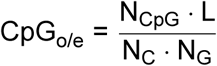

where N_CpG_ is the number of CpG dinucleotides, L is the sequence length, and N_C_ and N_G_ are the counts of C and G nucleotides, respectively. For analyses with *FAS* exon 6, we measured CpG composition as the number of CpG dinucleotides because of the more limited compositional diversity. For all experimental assays with measured exon inclusion levels, we computed model prediction error as the model prediction minus measured inclusion level (PSI). We plotted the effect of CpG composition on prediction error after binning sequences by PSI in bins of width 20% and grouping by CpG_o/e_. For each bin, we computed the median prediction error and interquartile range. Points with values outside 1.5*×* the interquartile range were removed during plotting as outliers. We tested for association between CpG composition and model prediction error, after controlling for PSI, using analysis of covariance (ANCOVA). Binning was used only for visualization; statistical tests were performed on unbinned data. We repeated the analysis using GpC composition (computed similarly to CpG) as a control.

To compute the enrichment of false positive SpliceAI predictions within genomic CpG islands, we first obtained a list of deep intronic CpG islands (at least 300 base pairs away from any exons annotated in any protein-coding transcripts). We identified false positive SpliceAI predictions as positions within all protein-coding genes with SpliceAI scores > 0.2 that were not annotated as splice sites in any protein-coding Ensembl transcript. For each CpG island, we computed the number of false positive predictions within 300 base pairs. We also selected an intronic window for comparison as a matched control from either the same gene or chromosome with the same length, a similar GC content (±3%), and not overlapping any other CpG islands. We computed the observed proportion of false predictions in CpG islands as well as the expected proportion using the matched comparison controls out of all false positive predictions. We also calculated top-k accuracy for all protein-coding genes with CpG islands before and after removing false positive predictions in intronic CpG islands. We obtained SpliceAI’s reported top-k accuracy on protein-coding transcripts and long non-coding RNA transcripts from the original publication.^4^

### CpG Methylation Analysis

To test whether CpG methylation is associated with changes in exon inclusion, we analyzed data from the DNMT1-inhibitor study by Pappalardi et al. 2021 (GEO number GSE135207).^17^ We downloaded whole-genome methylation measurements (from Illumina Methylation EPIC BeadChip arrays) and RNA-seq data from two matched samples of the MV4-11 leukemia cell line: MV411.

400nM.GSK032.d4.n2 (the time point and dose with maximal methylation inhibition) and MV411. 0nM.DMSO.d4.n2 (DMSO control from the same time point). All sequencing data were reprocessed using the same computational pipeline described in the original study. Briefly, RNA-seq reads were adapter-trimmed using fastp (v1.0.1),^38^ aligned to GRCh38 using STAR (v2.7.11b)^39^ with default parameters, and differential exon inclusion events were quantified using rMATS-turbo (v4.3.0).^40^

For each exon, methylation levels were computed as the mean methylation probe value across all probes overlapping that exon. Exon-level methylation depletion was defined as the log_2_ fold change in methylation between DNMT1-inhibitor-treated and DMSO samples. We then tested for associations between DNA methylation changes and differential exon inclusion by fitting a simple linear regression model, reporting the Pearson correlation coefficient and two-sided p-value. We also performed an analogous regression between exon CG count and differential exon inclusion.

### Stop Codon Analysis

For each sequence, we counted the number of stop codons (TAA, TAG, TGA) in any of the three frames within the putative exon. For all experimental assays with measured exon inclusion levels, we computed model prediction error as the model prediction minus measured inclusion level (PSI). We plotted the effect of stop codon count on prediction error after binning sequences by PSI in bins of width 20% and grouping by stop codon count. For each bin, we computed the median prediction error and interquartile range. Points with values outside 1.5*×* the interquartile range were removed during plotting as outliers. We tested for association between stop codon count and model prediction error using analysis of covariance (ANCOVA), controlling for PSI. Binning was used only for visualization; statistical tests were performed on unbinned data. We repeated the same analyses for each individual stop codon type.

For analysis of genomic predictions, we examined predicted 5’ and 3’ splice site scores of all internal protein-coding exons belonging to MANE Select transcripts. We only considered internal exons (i.e., not the first or last exon), as SpliceAI does not predict terminal exon boundaries. We compared splice site scores for final internal coding exons (i.e., exons that contain a stop codon marking the end of the coding sequence) with those for other internal coding exons. To control for exon length and rank, we also compared scores after binning exons by length and their position in the transcript. To control for the possibility of alternative splicing, we also compared scores of exons annotated as constitutive in Ensembl. For all comparisons, we used two-tailed Welch’s t-test to test for a difference between splice site scores of the two populations.

### RNA Secondary Structure Analysis

For sequences from the Liao et al. 2023 study,^18^ we predicted minimum free energy (MFE) RNA secondary structures using RNAfold from the ViennaRNA package^30^ (version 2.7.0) with default parameters. For input, we used each 70-nucleotide variable region surrounded by 10-nucleotide flanking upstream and downstream regions. We computed model prediction error as the model prediction minus measured inclusion level (PSI). We plotted the effect of predicted MFE on prediction error after binning sequences by PSI in bins of width 20% and grouping by predicted MFE. For each bin, we computed the median prediction error and interquartile range. Points with values outside 1.5*×* the interquartile range were removed during plotting as outliers. We tested for association between predicted MFE and model prediction error, after controlling for PSI, using analysis of covariance (ANCOVA). Binning was used only for visualization; statistical tests were performed on unbinned data.

For *MAPT* exon 10 variants, we used experimentally measured structural stability scores provided by Kumar et al. 2022.^27^ We computed model prediction error as the model prediction minus PSI. We regressed prediction error against structural stability scores and report the Pearson correlation coefficient. For *BLTP1*, we predicted minimum free energy RNA secondary structures using RNAfold with default parameters. Structures were plotted using VARNA version 3-93.^41^

### Genomic Sequences and Annotations

All analyses were conducted using human reference genome assembly GRCh38.p14. CpG island annotations (version 2022-10-18) were obtained from the UCSC Genome Browser. Gene and transcript annotations were obtained from Ensembl release 114. For analyses of protein-coding genes, we examined only transcripts tagged as MANE Select, except for our analysis of false positive splicing predictions, where we considered all Ensembl annotated protein-coding transcripts. For analyses of long non-coding RNA genes, we examined only transcripts tagged as Ensembl Canonical. For analysis of CpG methylation, we obtained chromosomal gene and transcript annotations from GENCODE Basic 49.

### Data Availability

All experimental datasets are publicly available from the cited sources. Trained model weights and processed predictions are available at https://github.com/regev-lab/splicing-interpretable-distillation.

### Code Availability

All code for model training, analysis, and figure generation is available at https://github.comregev-lab/splicing-interpretable-distillation.

## Acknowledgements

We thank Susan E. Liao, Arush Ramteke, and other members of the Regev Lab for comments. This work was supported by NSF MCB-2226731, NSF MCB-2246531, and a Simons Investigator Award. This work was also supported in part through NYU IT High Performance Computing resources, services, and staff expertise.

## Supplementary Information

**Figure S1.**
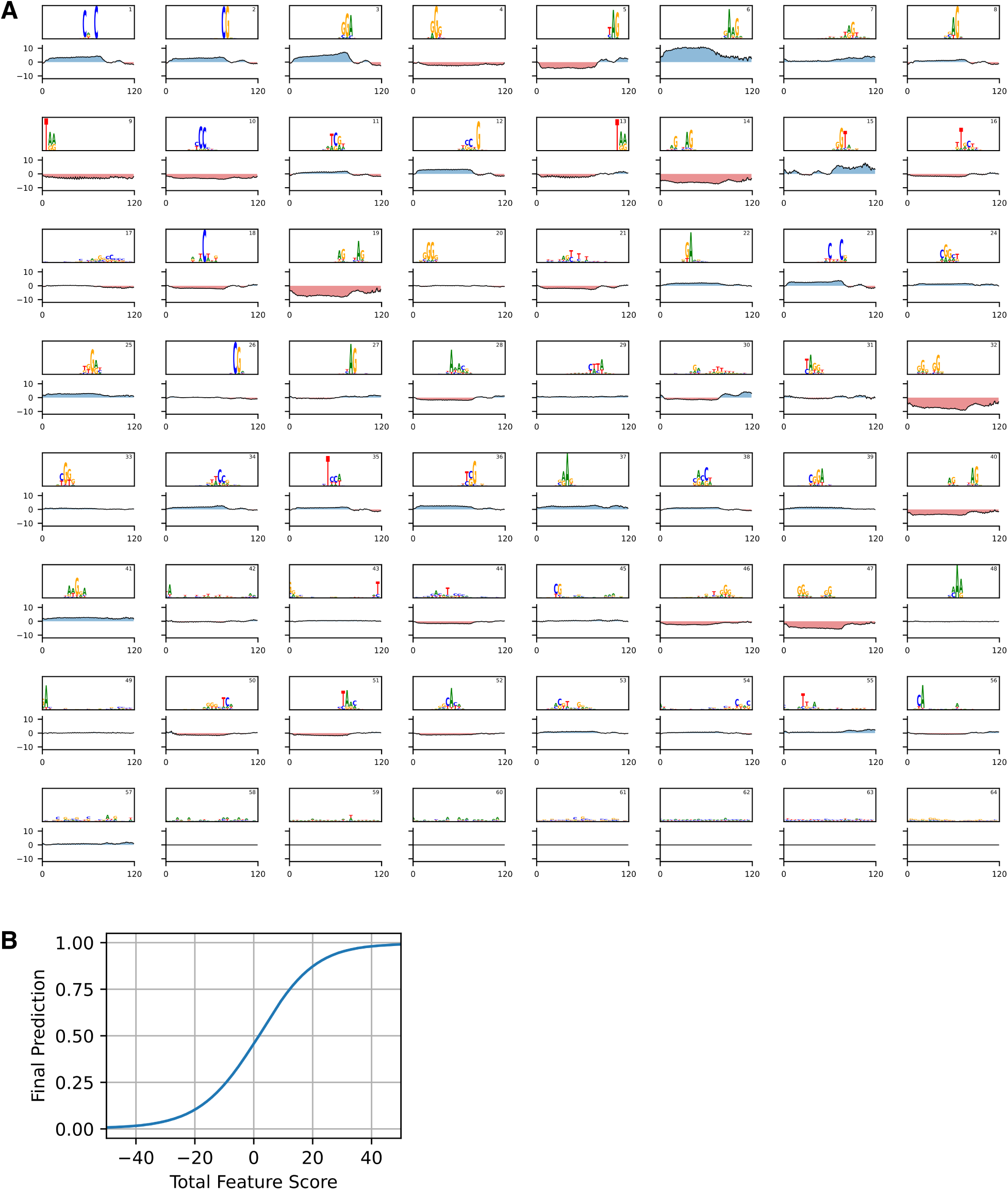
Interpretable distillation reveals exon recognition logic of SpliceAI (related to. **Figure 1). (A)** Sequence features learned by distilled SpliceAI model, sorted by importance rank (top right number). Sequence logos show convolutional kernel preferences. Positional score plots show position-dependent contributions along the 120 nt input region (70 nt variable region with 25 nt constant flanks). Blue: increases prediction; red: decreases prediction. **(B)** Learned transformation function mapping summed contribution scores to final predictions.

**Figure S2.**
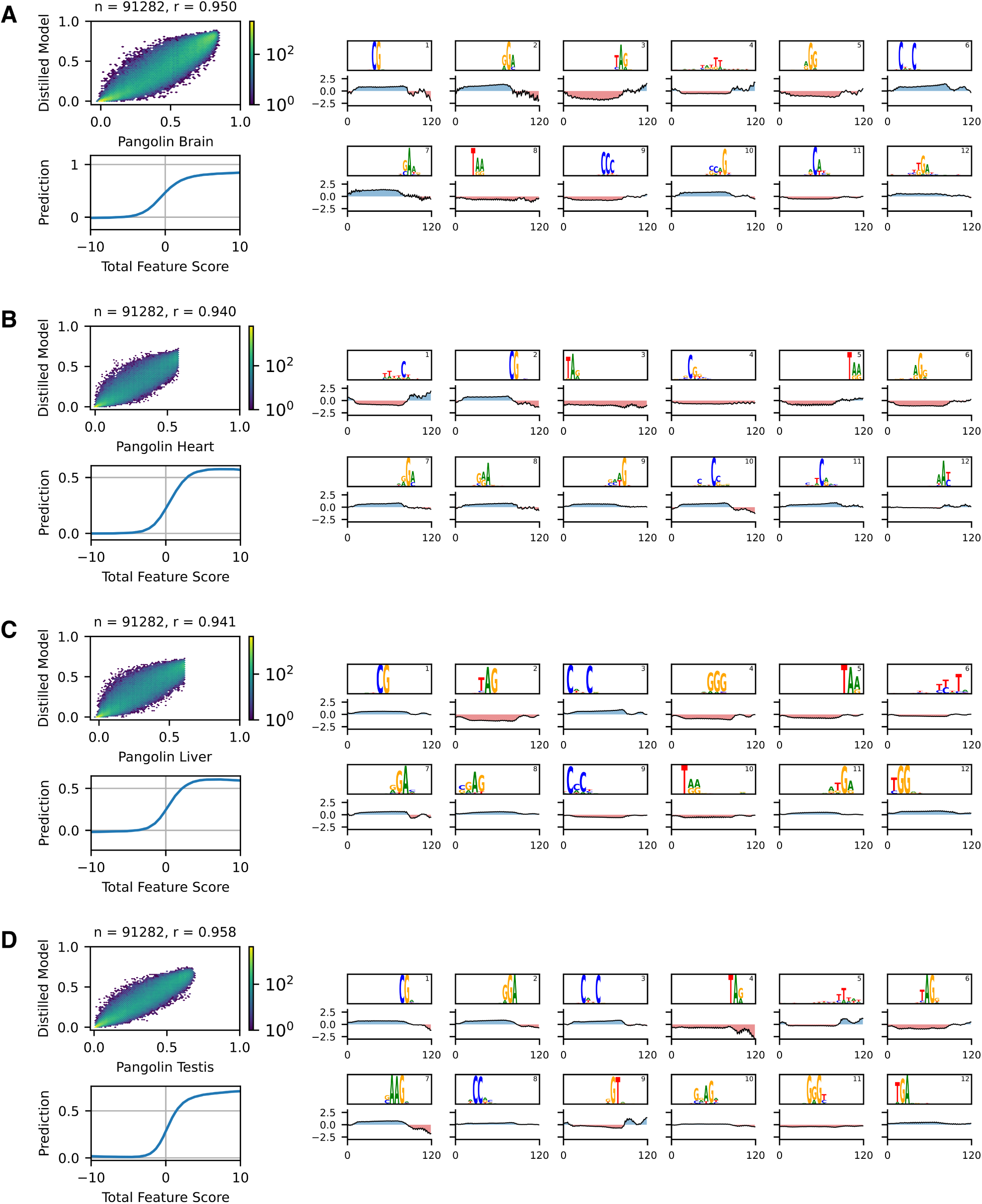
Interpretable distillation reveals exon recognition logic of Pangolin models (related to. **Figure 1). (A)** Pangolin brain model: (Top left) Comparison of distilled model predictions to Pangolin brain model predictions on held-out sequences. (Right) Top twelve sequence features learned by distilled Pangolin brain model, sorted by importance rank (top right number). Sequence logos show convolutional kernel preferences. Positional score plots show position-dependent contributions along the 120 nt input region (70 nt variable region with 25 nt constant flanks). Blue: increases prediction; red: decreases prediction. (Bottom left) Learned transformation function mapping summed contribution scores to final brain predictions. **(B)** Pangolin heart model, same format as (A). **(C)** Pangolin liver model, same format as (A). **(D)** Pangolin testis model, same format as (A).

**Figure S3.**
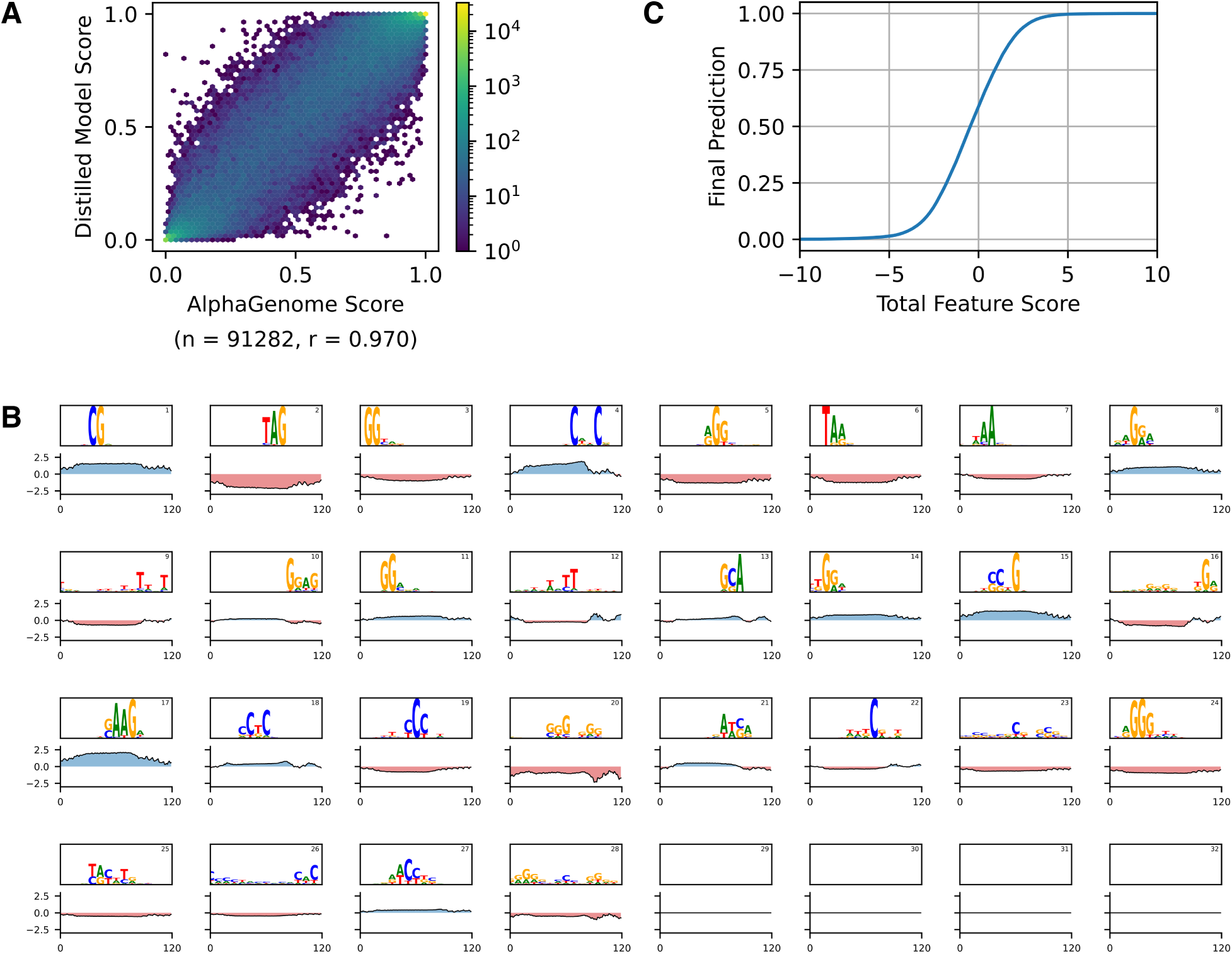
Interpretable distillation reveals exon recognition logic of AlphaGenome (related to. **Figure 1). (A)** Comparison of distilled model predictions to AlphaGenome predictions on held-out sequences. **(B)** Sequence features learned by distilled AlphaGenome model, sorted by importance rank (top right number). Sequence logos show convolutional kernel preferences. Positional score plots show position-dependent contributions along the 120 nt input region (70 nt variable region with 25 nt constant flanks). Blue: increases prediction; red: decreases prediction. **(C)** Learned transformation function mapping summed contribution scores to final predictions.

**Figure S4.**
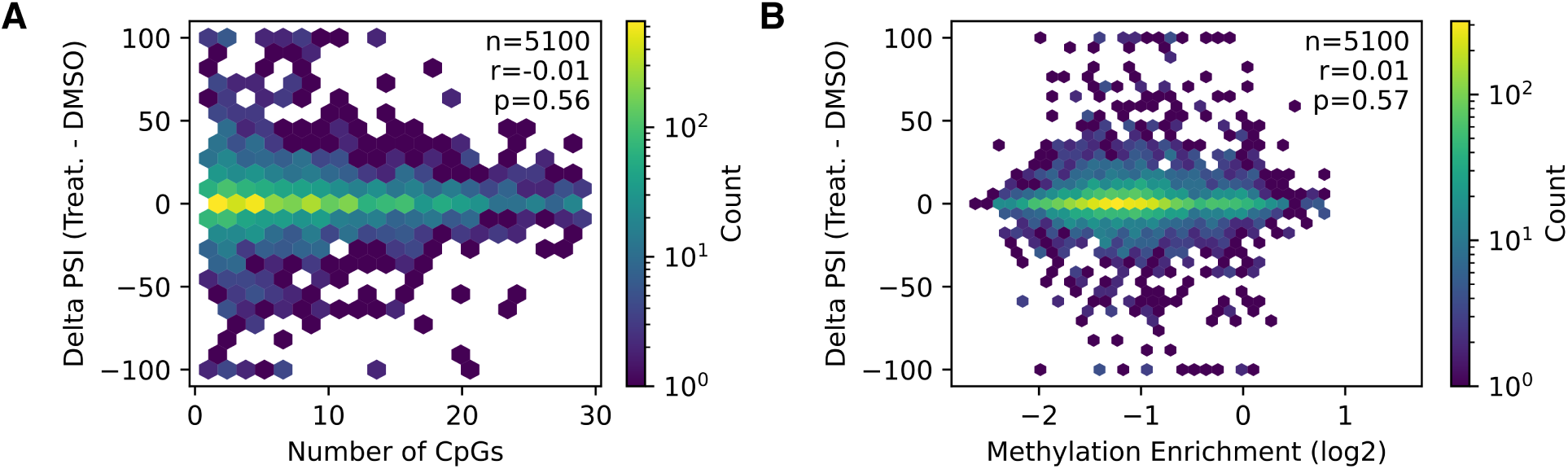
Analysis of CpG methylation and differential exon inclusion fails to reveal a significant association. **(A)** Number of CpGs per exon and differential exon inclusion (ΔPSI) following DNMT1 inhibitor treatment^17^ compared to DMSO control. **(B)** CpG methylation depletion per exon (relative to DMSO) and differential exon inclusion (ΔPSI).

**Figure S5.**
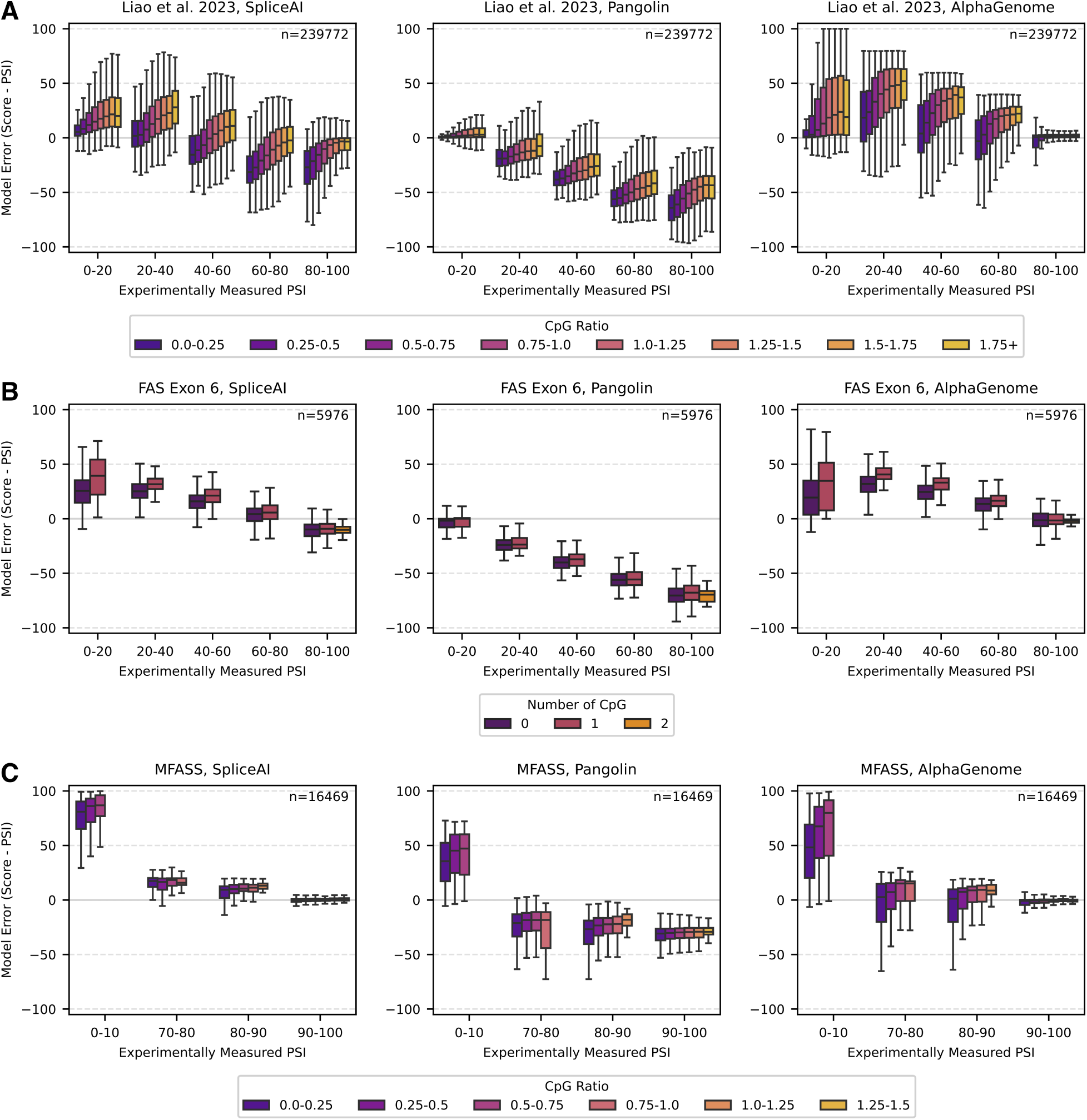
CpG enrichment confounds splicing predictions across experimental datasets (related to. **Figure 2A). (A)** Model prediction error (prediction minus measured inclusion) for SpliceAI, Pangolin, and AlphaGenome on Liao et al. 2023 library.^18^ Box plots show error binned by inclusion level (PSI) and grouped by CpG ratio. **(B)** Model prediction error on *FAS* exon 6 mutagenesis library.^19^ Box plots grouped by number of CpG dinucleotides. **(C)** Model prediction error on MFASS library.^20^ Box plots grouped by CpG ratio. Boxes in (A), (B), and (C) show quartiles; whiskers extend to the full distribution excluding outliers. Each box plot contains *≥*20 sequences.

**Figure S6.**
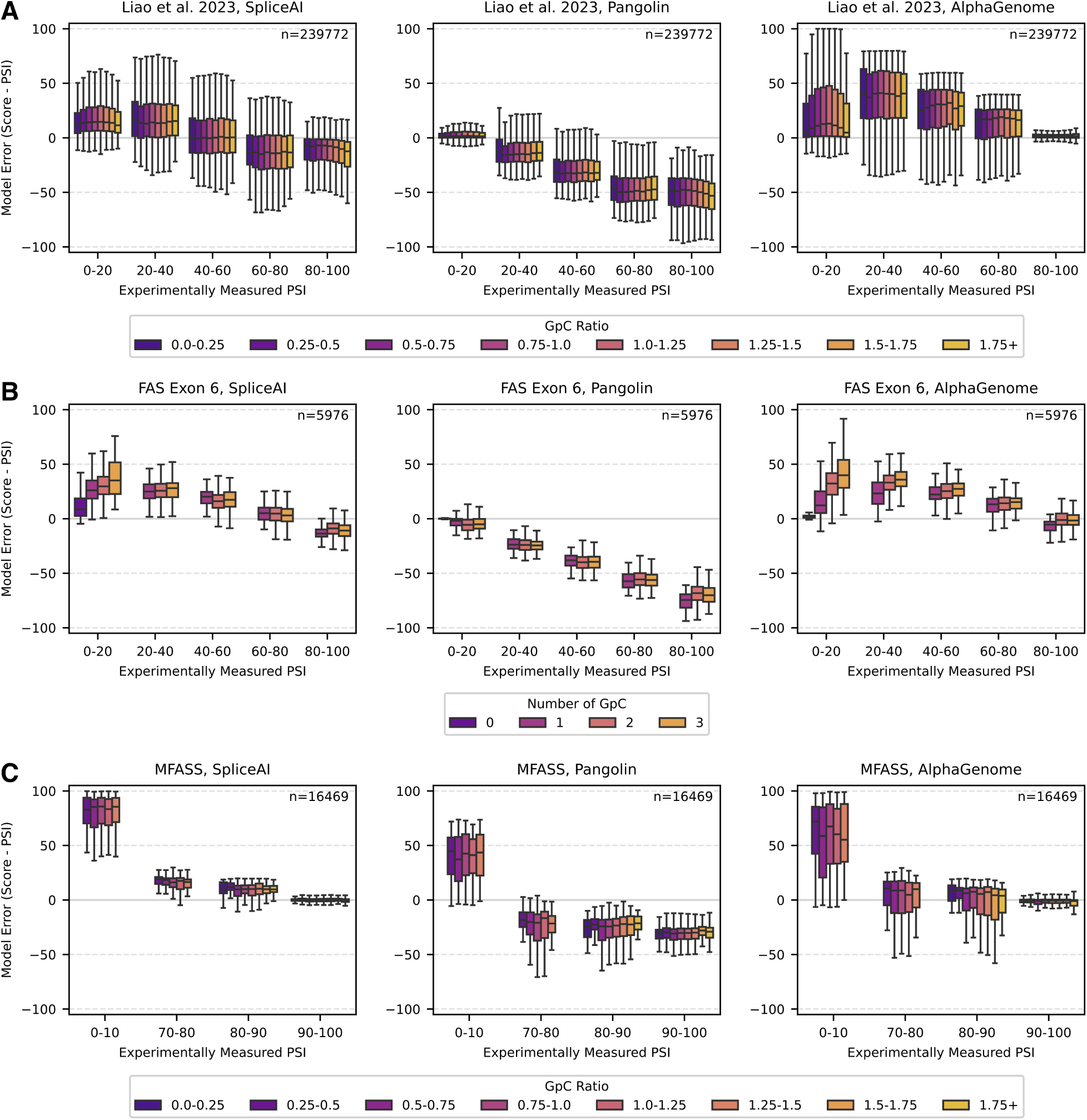
GpC enrichment does not systematically confound splicing predictions. **(A)** Model prediction error (prediction minus measured inclusion) for SpliceAI, Pangolin, and AlphaGenome on Liao et al. 2023 library.^18^ Box plots show error binned by inclusion level (PSI) and grouped by GpC ratio. **(B)** Model prediction error on *FAS* exon 6 mutagenesis library.^19^ Box plots grouped by number of GpC dinucleotides. **(C)** Model prediction error on MFASS library.^20^ Box plots grouped by GpC ratio. Boxes in (A), (B), and (C) show quartiles; whiskers extend to the full distribution excluding outliers. Each group contains *≥*20 sequences.

**Figure S7.**
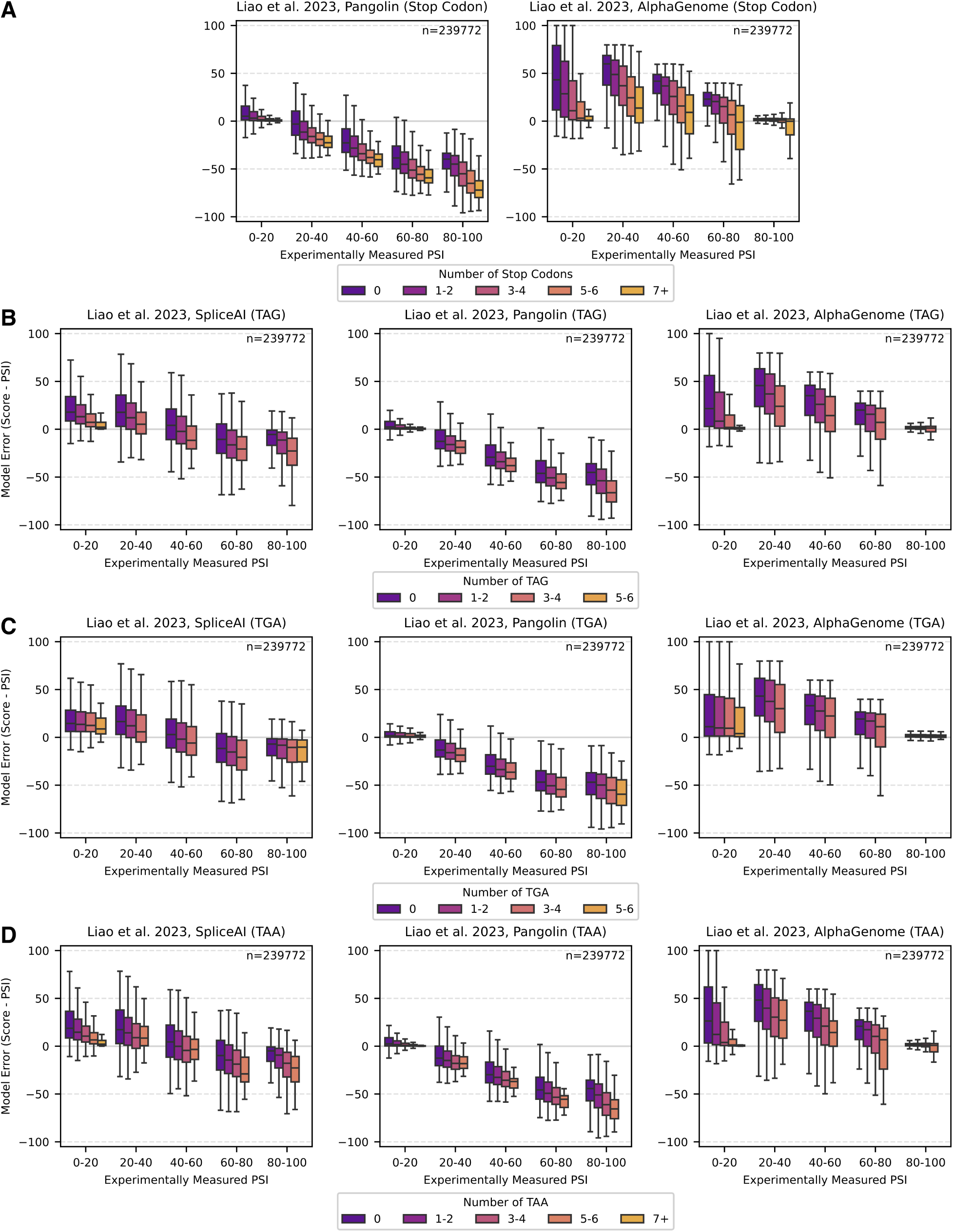
Stop codon presence confounds splicing predictions (related to. **Figure 3A). (A)** Model prediction error (prediction minus measured inclusion) for SpliceAI, Pangolin, and AlphaGenome on Liao et al. 2023 library.^18^ Box plots show error binned by inclusion level (PSI) and grouped by total stop codon count. **(B)** Error grouped by TAG codon count. **(C)** Error grouped by TGA codon count. **(D)** Error grouped by TAA codon count. Boxes in (A), (B), (C), and (D) show quartiles; whiskers extend to the full distribution excluding outliers. Each group contains *≥*20 sequences.

**Figure S8.**
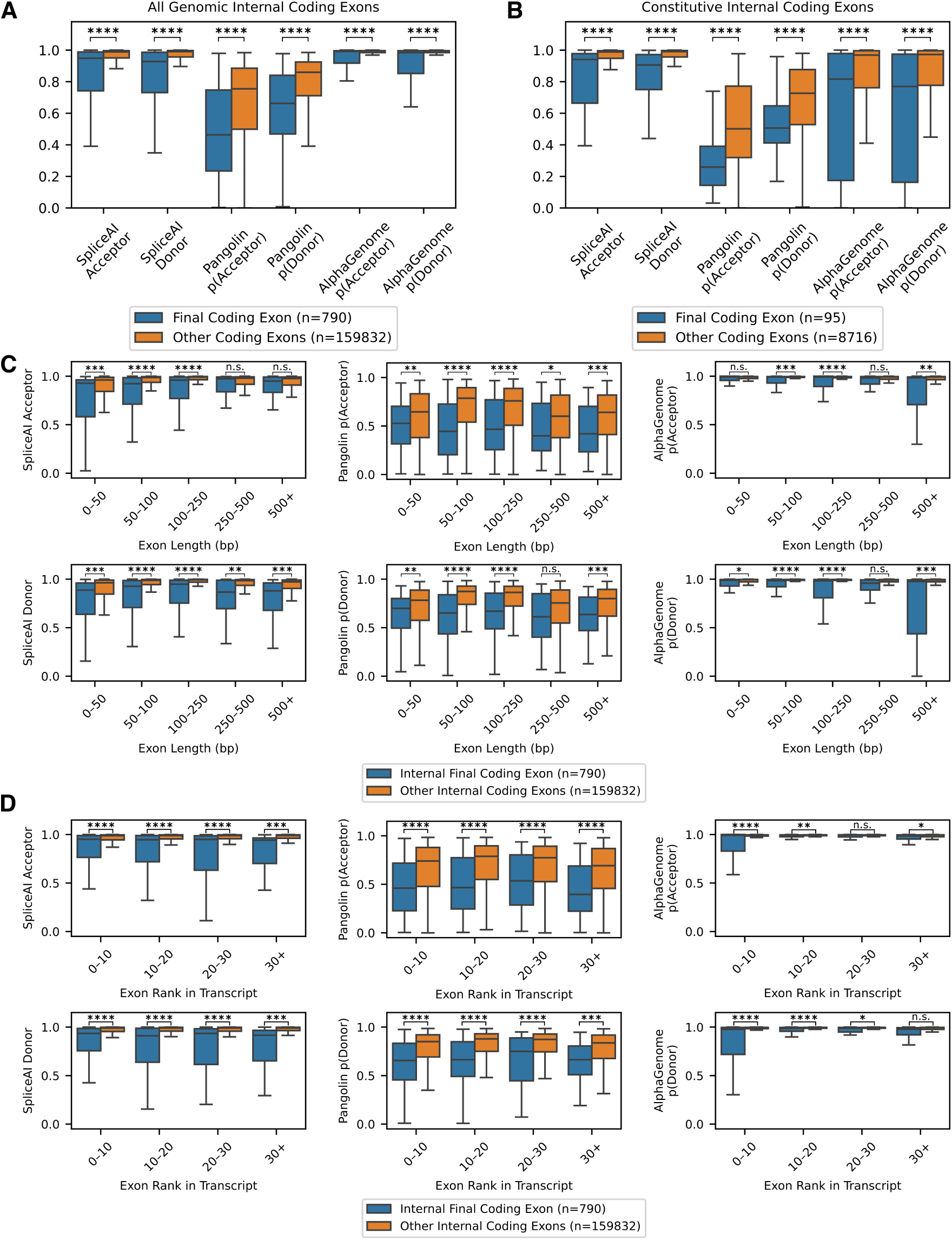
Stop codon-containing genomic exons receive lower splicing predictions (related to. **Figure 3B). (A)** Model predictions for splice acceptors and donors in final internal coding exons compared to in all other internal coding exons in MANE Select transcripts. **(B)** Model predictions for splice acceptors and donors in final constitutive internal coding exons compared to in all other constitutive internal coding exons in MANE Select transcripts. **(C)** Model predictions for splice acceptors and donors in final compared to in other internal coding exons, binned by exon length. **(D)** Model predictions for splice acceptors and donors in final compared to in other internal coding exons, binned by exon rank in transcript. Boxes in (A), (B), (C), and (D) show quartiles; whiskers extend to the full distribution excluding outliers. Two-tailed Welch’s t-test: ns: P > 0.05; *: P *≤* 0.05; **: P *≤* 0.01; ***: P *≤* 0.001; ****: P *≤* 0.0001.

**Figure S9.**
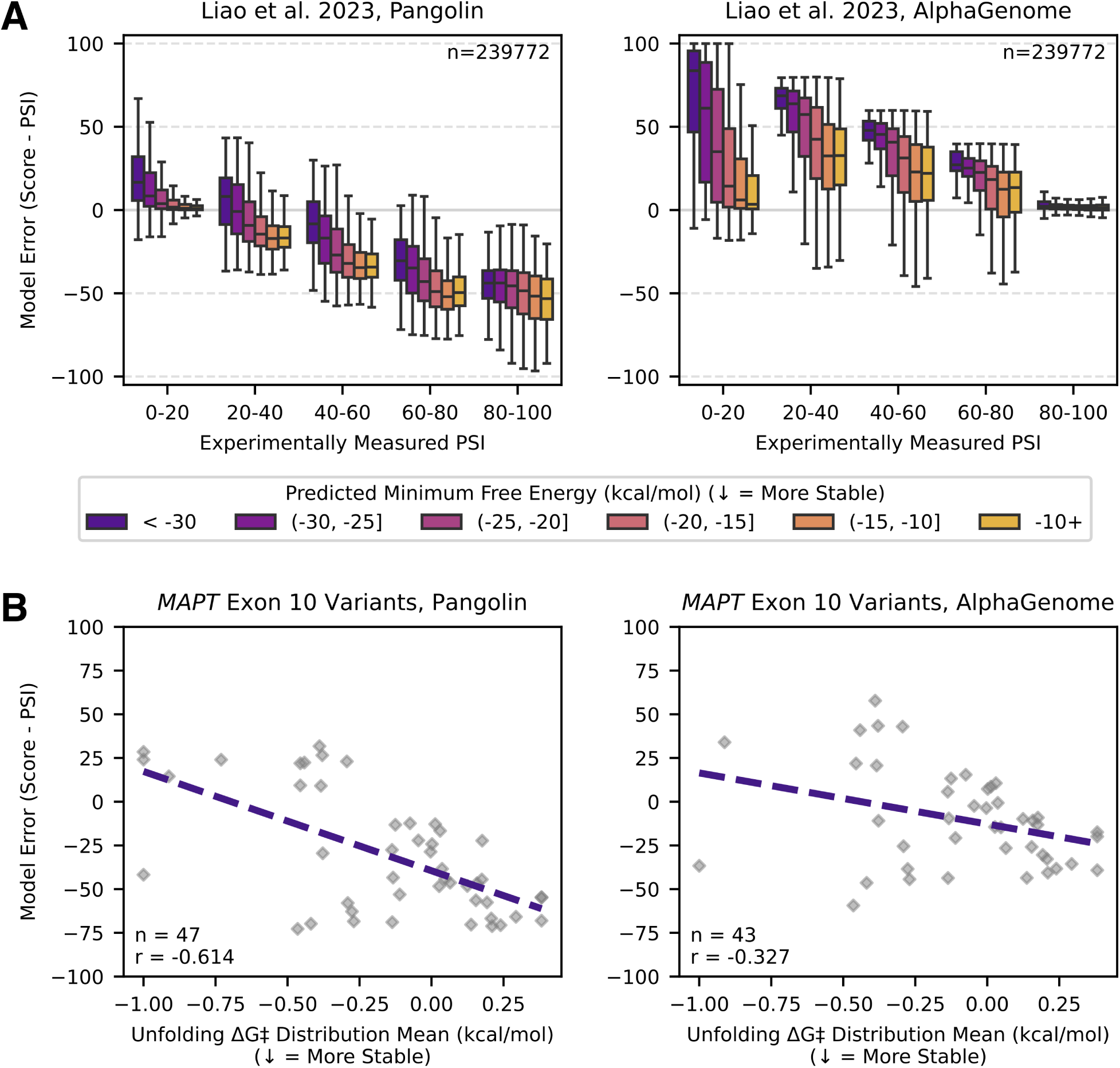
Splicing model predictions are blind to RNA structure (related to. **Figures 4A, 4B). (A)** Model prediction error (prediction minus measured inclusion) for Pangolin and AlphaGenome on Liao et al. 2023 library.^18^ Box plots show error binned by measured inclusion level (PSI) and grouped by predicted minimum free energy (MFE; more negative values indicate more stable structures). Boxes show quartiles; whiskers extend to the full distribution excluding outliers. Each box plot contains *≥* 90 sequences. **(B)** Model prediction error as a function of RNA structure score for Pangolin and AlphaGenome on *MAPT* exon 10 variants with experimentally characterized structures. Pearson correlation coefficient noted in bottom left. Four variants for which AlphaGenome failed to predict any splice junctions were excluded from AlphaGenome-specific analyses.

**Table S1.**
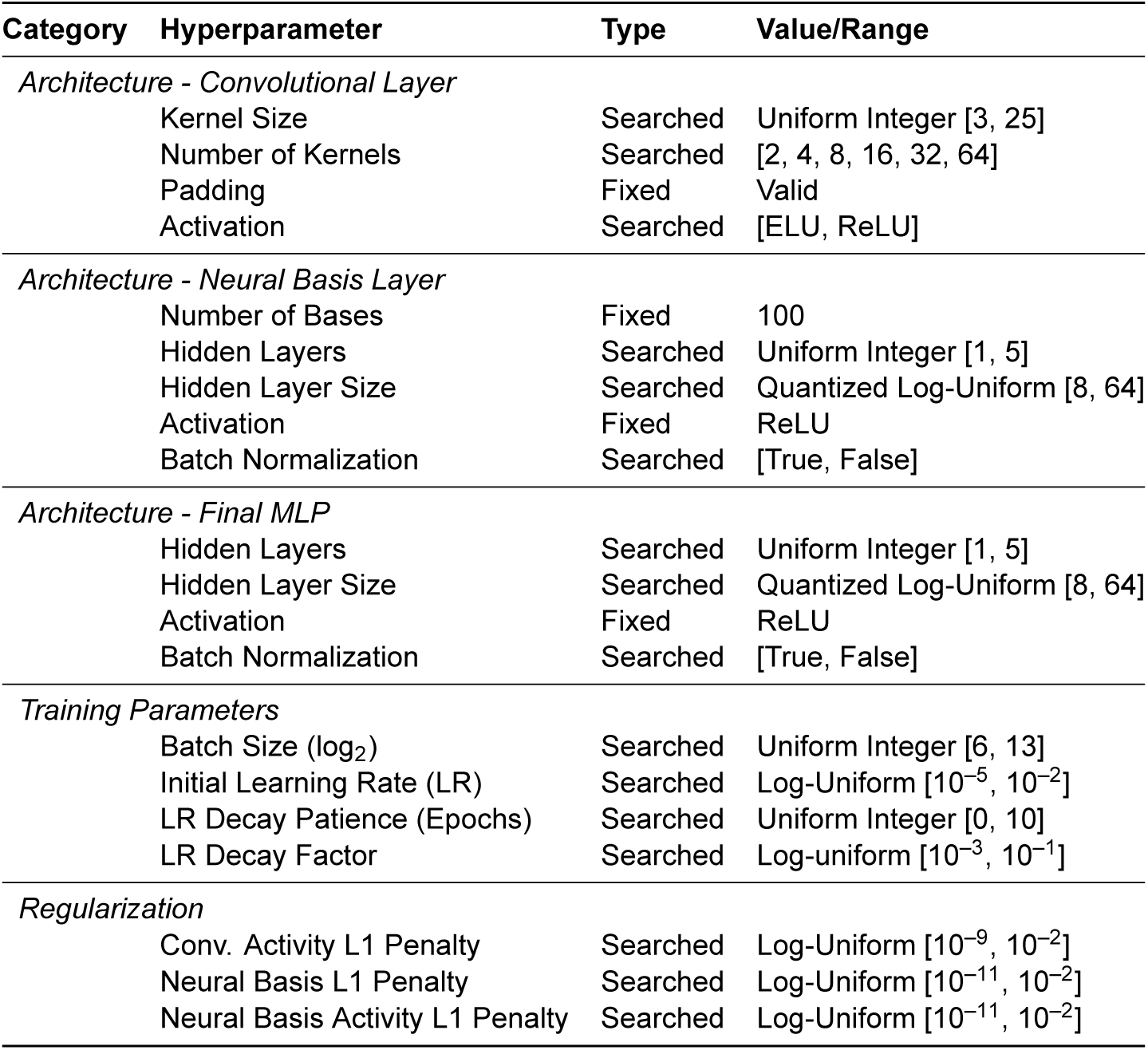
Hyperparameter search space for interpretable distillation models. Searched hyperparameters show the distribution and range used for optimization. Fixed hyperparameters show constant values used across all models. Conv.: convolutional; LR: learning rate.

| Category | Hyperparameter | Type | Value/Range |
| --- | --- | --- | --- |
| *Architecture - Convolutional Layer* |  |  |  |
|  | Kernel Size | Searched | Uniform Integer [3, 25] |
|  | Number of Kernels | Searched | [2, 4, 8, 16, 32, 64] |
|  | Padding | Fixed | Valid |
|  | Activation | Searched | [ELU, ReLU] |
| *Architecture - Neural Basis Layer* |  |  |  |
|  | Number of Bases | Fixed | 100 |
|  | Hidden Layers | Searched | Uniform Integer [1, 5] |
|  | Hidden Layer Size | Searched | Quantized Log-Uniform [8, 64] |
|  | Activation | Fixed | ReLU |
|  | Batch Normalization | Searched | [True, False] |
| *Architecture - Final MLP* |  |  |  |
|  | Hidden Layers | Searched | Uniform Integer [1, 5] |
|  | Hidden Layer Size | Searched | Quantized Log-Uniform [8, 64] |
|  | Activation | Fixed | ReLU |
|  | Batch Normalization | Searched | [True, False] |
| *Training Parameters* |  |  |  |
| | Batch Size ($\log_2$) | Searched | Uniform Integer [6, 13] |
| | Initial Learning Rate (LR) | Searched | Log-Uniform [$10^{-5}$, $10^{-2}$] |
|  | LR Decay Patience (Epochs) | Searched | Uniform Integer [0, 10] |
| | LR Decay Factor | Searched | Log-uniform [$10^{-3}$, $10^{-1}$] |
| *Regularization* |  |  |  |
| | Conv. Activity L1 Penalty | Searched | Log-Uniform [$10^{-9}$, $10^{-2}$] |
| | Neural Basis L1 Penalty | Searched | Log-Uniform [$10^{-11}$, $10^{-2}$] |
| | Neural Basis Activity L1 Penalty | Searched | Log-Uniform [$10^{-11}$, $10^{-2}$] |

